# Ultra-high dose oral *ω*3 docosahexaenoic acid (DHA) or eicosapentaenoic acid (EPA) block tumorigenesis in a *MYCN*-driven neuroblastoma model

**DOI:** 10.1101/2024.05.30.596731

**Authors:** Vishwa Patel, Yan Ning Li, Lorraine-Rana E. Benhamou, Hui Gyu Park, Mariya Raleigh, J. Thomas Brenna, John T. Powers

## Abstract

**Background/Objectives:** Neuroblastoma is a genetically diverse, highly metastatic pediatric cancer accounting for 15% of childhood cancer deaths despite only having ~8% of childhood cancer incidence. The current standard of care for high-risk disease is highly genotoxic. This, combined with less than 50% survival in high-risk disease and an abysmal 5% survival in relapsed cases, makes discovering novel, effective, and less toxic treatments essential.

**Methods:** A prophylactic syngeneic mouse model was used to test high-dose lipid-mediator highly unsaturated fatty acids on tumorigenesis. Wild-type mice were gavaged with 12.3-14.6 g/d (adult human equivalent) omega-3 EPA, DHA, or oxidation-resistant bis allylic deuterated DHA (D-DHA) and 4.6-6.0 g/d arachidonic acid (ARA). At seven days, *MYCN*-expressing murine neuro-2a cells syngeneic to the gavaged mice were injected subcutaneously. Oral gavage continued for 10-20 d post-injection when tumors and tissues were harvested.

**Results:** Fifty percent of control (not gavaged) animals form tumors (4/8) at about 10 d. High-dose DHA, D-DHA, and EPA block tumor formation completely in n=8 or 10 animals. In contrast, ω6 arachidonic acid (4.6-6.0 g/d) enhances tumor formation (6/10 tumors) and reduces latency (5.5 to 10 days)compared to control. Co-delivery of ARA and EPA results in a reduced tumor burden analogous to the control group, suggesting that EPA directly opposes the mechanism of ARA-mediated tumor formation. DHA acts through a non-oxidative mechanism.

**Conclusions:** Sustained high dose ω3 (weeks/months) is safe and well tolerated in humans. These results suggest that ω3 DHA and EPA delivery at ultra-high doses may represent a viable low-toxicity therapy for neuroblastoma.

**Simple Summary:** Pediatric Neuroblastoma has an overall mortality rate above 50%, and the current standard of care consists of highly genotoxic compounds. The biological actions of omega-6 (ω6) and omega-3 (ω3) highly unsaturated fatty acids (HUFA) generally oppose one another with the ω6 HUFA signaling for inflammation and angiogenesis (new blood vessel formation). Prolonged use of ultrahigh dose (15-20 g/d) ω3 HUFA has shown efficacy in catastrophic human traumatic brain injury and is well tolerated. Tumors form in about 50% of mice in our pediatric neuroblastoma model. We show that 12-14 g/d adult human equivalent doses of ω3 EPA or DHA, as well as an oxidation-resistant form of DHA (D-DHA), completely block tumor formation, whereas a dose of about 5 g/d of ω6 ARA enhances tumorigenesis. Our data suggest that ultra-high dose ω3 therapy should be carefully investigated as a low-toxicity approach to neuroblastoma intervention.

## 1. Introduction

Neuroblastoma (NB) is a highly metastatic pediatric cancer that accounts for 10-15% of childhood cancer deaths [1,2]. At diagnosis, about 40% of patients present with high-risk (HR) disease. Unlike low- and intermediate-grade NB, where survival has improved over time, HR and recurrent NB retain less than 50% and 5% survival rates, respectively. The extensive treatment regimen for HR NB is also a cause for concern. The current standard of care (SoC) for these patients includes surgical resection, high-dose combination chemotherapy, and radiation [3]. Post-NB treatment, many children are left with lingering, life-long issues affecting the cardiovascular, endocrine, and excretory systems, as well as an increased incidence of secondary malignancies later in life. Therefore, novel, less toxic therapies are needed to treat HR and relapsed neuroblastoma.

Approximately 25% of NB cases exhibit amplification of the proto-oncogenic transcription factor *MYCN* [3,4]. *MYCN* prevents neuronal differentiation, instead promoting cell proliferation and apoptotic resistance[3,5]. It has also been shown to alter cellular metabolism by increasing dependence on glutamine and fatty acid uptake [3,6]. Fatty acids themselves play multiple roles within the cell. They are an essential structural component of phospholipid membranes, making their synthesis or uptake integral to cell growth. Highly unsaturated fatty acids (HUFA), polyunsaturated fatty acids with more than three double bonds, can be released from cell membranes by enzymes from the PLA2 family and converted to eicosanoids and docosanoids. These compounds, collectively known as oxylipins, serve a fundamental role in paracrine signaling and include immune and angiogenesis-stimulating species such as prostaglandins, 20-HETE, and leukotrienes [7]. Although fatty acids can contribute to tumor formation or inhibition depending on the cancer type and mutations present [8–10], in NB, UFAS gene expression is strongly tied to patient outcome [11]. UFAS genes *FASN, ELOVL6, SCD, FADS2*, and *FADS1* are upregulated in HR NB and strongly correlate with decreased overall survival [11]. Transformation to malignancy was also found to alter fatty acid synthesis and levels within NB cells, further cementing that fatty acid content influences tumorigenesis both *in vitro* and *in vivo* [11].

HUFAs, both omega-3 and omega-6, can be derived endogenously from essential precursors α-linolenic and linoleic acid. HUFAs docosahexaenoic acid (DHA) and eicosapentaenoic acid (EPA), both ω3, as well as ω6 arachidonic acid (ARA), have been shown to inhibit cancer cell growth via the generation of reactive oxidative species (ROS) *in vitro* [12–15]. Human trials using between 1-2g/day ω3 HUFA have yielded mixed results [15]. High doses of ω3 fatty acids (up to 20 g/d), however, have been successful in treating patients with traumatic brain injury, hypertriglyceridemia, or hypertension and have shown that these treatments are well tolerated in humans [16–18]. High-dose ω3 HUFA has not previously been explored in cancer. Therefore, administration of these fatty acids may constitute a novel treatment approach to neuroblastoma.

Here, we aimed to understand the effects of high-dose ω3 and ω6 HUFA on tumor formation using dietary exogenous HUFA to modulate tumorigenesis in a syngeneic mouse model of *MYCN*-driven NB. Also, a deuterium-substituted, ROS-resistant isotopologue of DHA (*bis* allylic deuterated DHA; D-DHA) was used to investigate the impact of oxidation sensitivity on the mechanism of DHA [19,20]. Combined dietary ARA and EPA exposure was also evaluated.

## 2. Materials and Methods

### 2.1. Diets and oils

Mice were fed on customary facility mouse chow with a composition approximating AIN-93G [21]. ARA was from a single-cell triglyceride oil with about 44% ARA, with the rest composed of monounsaturated and saturated fatty acids (ARASCO, DSM, Columbia MD). EPA and DHA were from dietary supplement triglyceride oils of about 88% EPA and 82% DHA, respectively. The EPA and DHA supplements, in capsule form, were obtained from a vendor on Amazon.com and, according to labels, were purified from a mixture of natural fish oils. Oils were analyzed in-house before use, as outlined below. Doses and human equivalent doses are presented in ***Table 1***.

**Table 1.**
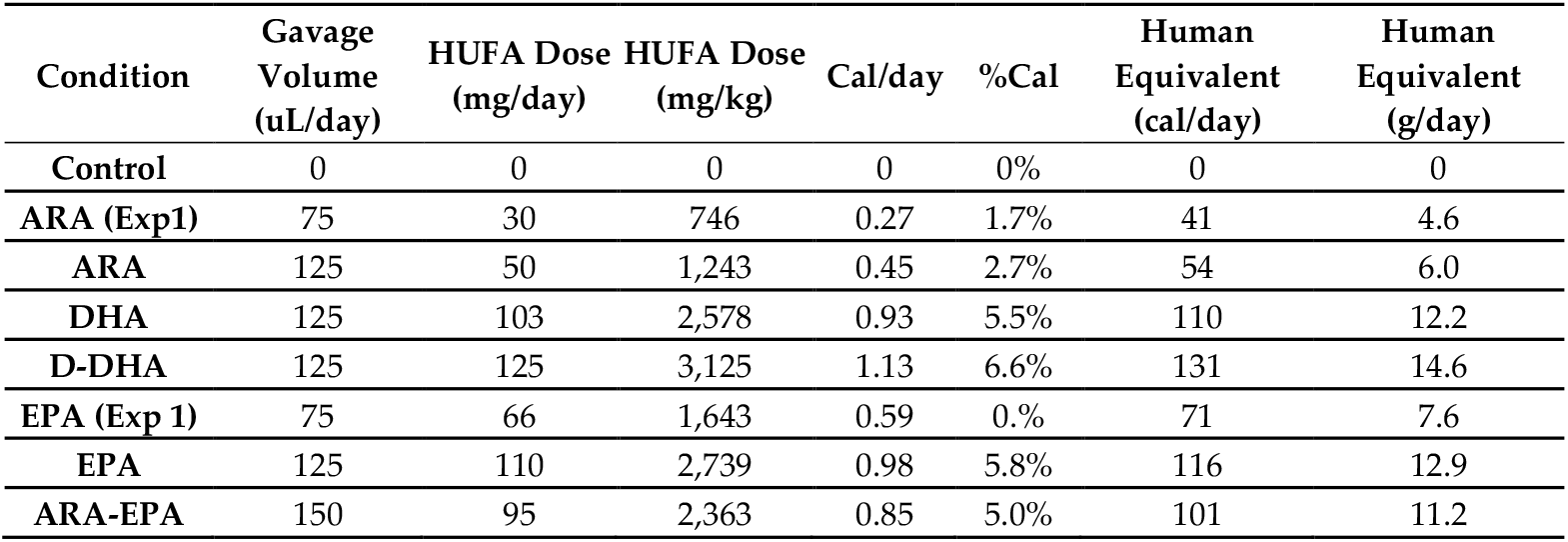
Dosing and caloric intake information for arachidonic acid (ARA), eicosapentaenoic acid (EPA), docosahexaenoic acid (DHA), and deuterated DHA (D-DHA) for our gavaging protocol, where mice were gavaged every other day. The table includes the highly unsaturated fatty acids (HUFA) dose in mg/day and mg/kg body weight/day. It shows the equivalent human dose assuming a daily intake of 2000 calories.

### 2.2. Cell Lines and Mice

Murine neuroblastoma Neuro-2a cells transduced with a human *MYCN* transgene were initially acquired from American Type Tissue Culture (ATCC-CCL131) and were cultured in RPMI-1640 media supplemented with 10% heat-inactivated fetal bovine serum (HI-FBS). Cells were grown at 37°C in a water-saturated atmosphere of 95% air and 5% CO2. Cells were tested monthly and consistently found to be negative for mycoplasma contamination. Neuro-2a cells are widely used as an *in vivo* neuroblastoma tumor model, typically in immune-compromised mice[22–25]. We established our Neuro-2:*MYCN* system [26] using syngeneic, wildtype strain A/J mice purchased from The Jackson Laboratory (Jaxmice strain #000646).

### 2.3. Plasmids

The *MYCN* pPB[Exp]-EF1A>EGFP(ns):P2A:hMYCN[NM_001293228.2 construct (de-posited at Addgene.org) is of our design and constructed by Vector Builder. Neuro-2a cells were reverse-stably transfected with the *MYCN* construct (Neuro-2a: *MYCN*), using lipofectamine™ 3000 transfection reagent (Invitrogen™) and a transposase plasmid (pRP[Exp]-mCherry-CAG>hyPBase) at a one µg concentration into six-well plates. Then, fluorescence-activated cell sorting (FACS) techniques were used to isolate transfected cell populations, and cells were used in further experiments.

### 2.4. Mouse oral gavage and injection procedures

#### 2.4.1. Experiment 1

A total of 8 mice per group were orally administered ARA (150 uL) and EPA (150 uL) for seven days before the injection of cells, respectively; on day 7, 2.5×10^6^ Neuro-2a:*MYCN* transfected cells were subcutaneously injected. For the ten days following cell injection, mice received oral administration of ARA and EPA by gavage every other day. At 10 days post-injection, the mice were euthanized, tumors were collected, and tumor weight and dimensions were measured. Tumor volume was calculated by using the formula *V* = 0.5 × *L* × *W*^*2*^, where *V* = tumor volume, *L* = Length, and *W* = tumor width.

#### 2.4.2. Experiment 2

A total of 10 mice per group were orally administered ARA (250 uL), EPA (250 uL), a combination of ARA/EPA (150 uL each), DHA (250uL), or D-DHA (250 uL) for seven days before the injection of cells, respectively, on day 7, 2 × 10^6^ Neuro-2a:*MYCN* transfected cells were subcutaneously injected into each mouse. For 20 days following cell injection, mice received oral ARA, EPA, ARA/EPA, DHA, or D-DHA every other day. Vernier calipers were used to measure tumor growth every other day until the study’s conclusion. At 20 days post-injection, mice were euthanized, final tumor measurements were gathered, and tissue/tumor samples were collected for fatty acid analysis; tumor volume was calculated as in experiment 1.

### 2.5. Fatty Acid Analysis

Doses and tissues were analyzed according to routine methods in our laboratory [27]. Briefly, tissue samples are minced to apparent homogeneity, or oils are used as is and treated with an aqueous phase to liberate fatty acids and convert them to fatty acid methyl esters (FAME) using previous methods [28]. Briefly, in one tube, an aqueous phase converts glycerolipid fatty acyl groups into fatty acid methyl esters (FAME), which enter an organic phase. FAME mixtures are evaporated, resuspended in heptane, and injected into a gas chromatograph (GC) equipped with a BPX-70 capillary column. An equal weight standard is used to verify response factors. Data are expressed as % weight-for-weight (% w/w) of total fatty acids.

### 2.6. RT-qPCR

Tumor tissues were harvested, minced, and homogenized using a mechanical homogenizer, followed by total RNA extraction with (TRIzol™, 15596026) Reagent according to the manufacturer’s protocol. RNA quality and integrity were assessed using Nanodrop. Genomic DNA contamination was eliminated through DNase treatment during RNA extraction. cDNA synthesis was performed using a Verso cDNA Synthesis Kit (Thermo Scientific™, catalog#AB1453A). Real-time qPCR was conducted using a SYBR Green detection system on a (Bio-Rad CFX96), with primers for *Fads1, Fads2, Elovl2*, and Elovl5 (all the primers acquired from IDT™). Each 20 µL reaction included SYBR Green master mix, primers (200 nM final concentration), and 1 µL of cDNA (Final concentration 1ng/µL). Relative gene expression was calculated using the 2^−ΔΔCt method, normalizing target genes to the endogenous control (GAPDH), validated for stability under experimental conditions. Primer sequences were as follows: *Fads1* forward CCACCAAGAA-TAAAGCGCTAAC, reverse AGCAGGTAGACCAGGAAGA; *Fads2* forward CATGAC-TATGGCCACCTTTCT, reverse GCTGAGGCACCCTTTAAGT; *Elovl2* forward ACATGTTTGGACCACGAGATT, reverse GTACGTGATGGTGAGGATGAAG; *Elovl5* forward CTATGAGTTGGTGACAGGTGTG, reverse TGGAGAAGTAGTACCAC-CAGAG. All data were analyzed as fold changes relative to controls (GAPDH), with mean ± standard deviation from biological and technical replicates, and statistical analysis was performed using GraphPad Prism, Version 3.1.

### 2.7 Statistical Analysis

All statistical analyses were conducted using GraphPad Prism, Version 3.1. Data are presented as mean ± standard error of the mean (SEM), unless otherwise specified. Tumor volumes and other continuous variables were analyzed using one-way or two-way analysis of variance (ANOVA) followed by Tukey’s post-hoc test for multiple comparisons. The differences in tumor latency between groups were analyzed using a t-test, with *p<0.05 and **p<0.01. Fatty acid composition and gene expression data were analyzed by two-way ANOVA followed by Tukey’s multiple comparison test to assess the interaction between treatment groups and tissue type. A pp-value of < 0.05 was considered statistically significant. Statistical significance levels are: *p< 0.05, **p< 0.01,<0.001,<0.0001. Each figure in the manuscript includes specific statistical details relevant to the data presented, ensuring clarity regarding the methods used for each analysis.

### 2.8. Ethics

Mouse protocols used in this study were approved by the University of Texas IACUC committee (IACUC protocol: AUP-2023-00058).

### 3. Results

### 3.1. Mouse diet and gavage oils

The oils used for oral gavage were analyzed for fatty acid profile (***Supplemental Figure 1, 2***). The ARA, EPA, and DHA concentrations were 44.5 %,w/w, 87.6%,w/w, and 82.5%,w/w, respectively. Base diet had 0.13%,w/w ARA, 0.61%,w/w EPA, and 1.30%,w/w DHA incidental to the addition of fish meal to an otherwise conventional AIN-93G formula (***Supplemental Figure 3***). Cells expressing either GFP control or MYCN protein revealed 4.2%,w/w ARA, 0.85%,w/w EPA, and 0.6%,w/w DHA levels in Neuro-2a:*MYCN* cells, none of which were significantly different from GFP control (***Supplemental Figure 4***).

### 3.2. Dosing

Two experiments were performed sequentially (Table 1). Experiment 1 used Control (no gavage), ARA, and EPA, both gavaged at 75 uL oil per day. Experiment 2 had Control (no gavage), and ARA and EPA at a higher dose of 125 uL per day, thus enabling a dose comparison, and a combined dose of ARA and EPA at 75uL each, the same doses as experiment 1. Experiment 2 also used DHA and D-DHA at 125 uL. Adult human dose equivalents calculated based on caloric intake of 2,000 cal/d show that the two ARA doses were 4.6 and 6.0 g/d, EPA were 7.6 and 12.9 g/d, and DHA and D-DHA were 12.2 and 14.6 g/d, respectively.

### 3.3. ARA and EPA have opposing effects on tumor formation in a syngeneic neuroblastoma model

Wildtype strain A/J mice syngeneic with Neuro-2a cells were gavaged every other day for seven days before subcutaneous injection of 2.5×10e6 Neuro-2a:*MYCN* cells into the left flank, n=8 per group. Mice gavaged with 75ul/day ARA displayed a significantly accelerated tumor incidence, with an average tumor latency of 5.25 days (***Figure 1a***), compared to 7 days in sham-gavaged mice. In contrast, the EPA-gavaged mice group demonstrated significantly reduced tumor incidence, with only three tumors developing with an average latency of 10 days (***Figure 1a***). Comparing tumor volumes across the two groups, ARA-gavaged mice exhibited the highest tumor burden, with significantly larger tumors than the EPA-gavaged group (***Figure 1b, 1e, Supplement Figure 5a, 5c***).

**Figure 1.**
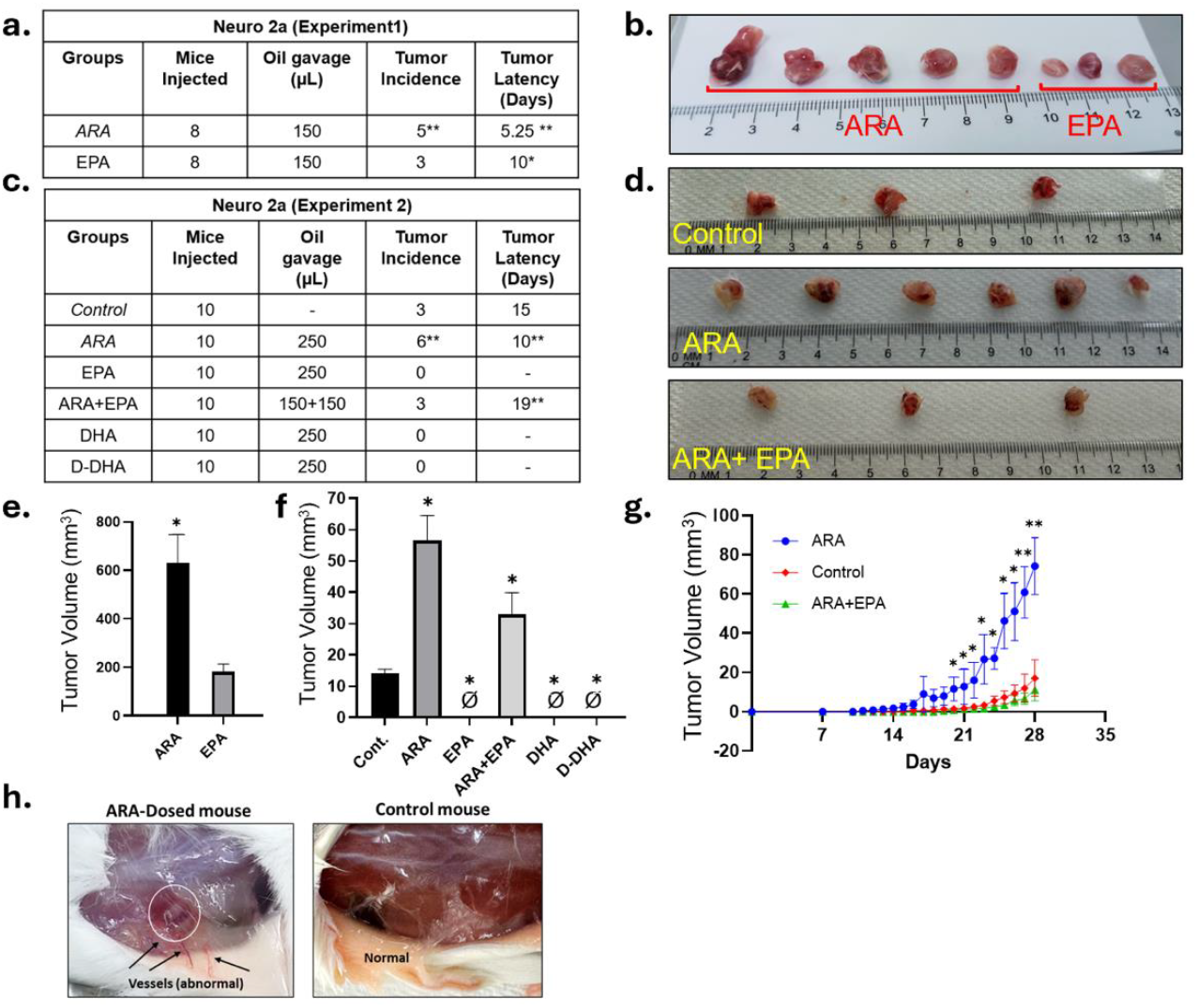
Syngeneic cell-derived xenograft (CDX) model of MYCN-driven neuroblastoma: ARA and EPA were administered orally every 48 hours to the mice for seven days before cell injection. 2.5*10^6 MYCN expressing Neuro-2a cells were injected subcutaneously into the mice. ARA, EPA, and ARA+EPA feeding continued until the study endpoint in both experiment rounds. a), c) Summary of tumor incidence and latency across groups treated with ARA, EPA, and control in Exp1 and 2, respectively, Significance was calculated using a t-test: *p<0.01 and **p<0.001, P<0.0001 b), d) Tumor images and relative sizes. e, f) A bar graph depicting the change in tumor volume across different experimental groups shows the highest tumor volume in the ARA-treated group. The slash symbol indicates that no tumors emerged in that condition (right panel). Significance was determined using a t-test, with *p<0.05 and **p<0.01 indicated. g) XY plot displaying the tumor volume changes across various treatment groups over time (experiment 2). Significance was calculated using a t-test: *p<0.05 and **p<0.01. h) Images of vessel formation in ARA-gavaged mice compared to control (experiment 2).

We then performed a second experiment with n=10 mice in each group. Doses were adjusted upward for both ARA and EPA, and a group with a combined dose of ARA-EPA. Similar to Experiment 1, ARA-gavaged mice again displayed significantly earlier tumor latency, increased tumor incidence, and tumor size than the control (***Figure 1c, d, f, Supplemental Figure 5b, 5d***). In addition, the ARA-gavaged mice developed visible robust new vasculature not seen in any other condition (***Figure 1h***). Mice gavaged with the ARA/EPA combination yielded tumors at the same incidence as control. However, their latency was significantly delayed, and their size was smaller as well, suggesting that EPA may be both beneficial in preventing tumor formation and dominant over pro-tumorigenic ARA effects (***Figure 1d, 1f, 1g, Supplemental Figure 5b, 5d***). Strikingly, the higher EPA (only) dose completely blocked tumor formation, yielding 0 tumors out of any of the 10 mice (***Figure 1c, 1f, 1g***).

### 3.4. Both DHA and D-DHA block tumor formation

Our only DHA- and D-DHA dosings were similar to the higher dose of EPA (12.2, 14.6 vs 12.9, Table 1). Similar to the higher EPA dose, gavaging DHA and D-DHA completely abolished tumor formation (***Figure 1c, 1f, 1g***).

### 3.5. Tumor and Tissue Fatty Acid Accumulation

We next analyzed emergent tumors and normal liver and skeletal muscle for changes in HUFA levels in response to oral gavage. Gavage ARA increased tumor and muscle ARA but not liver ARA (***Figure 2a,2b, left panels, Supplemental Figure 6***). Tissue EPA remained very low (below 1%) in tumors, liver, and skeletal muscle, even for EPA-gavaged animals (***Figure 2a, 2b, middle panels, Supplemental Figure 6***). DHA levels were also unchanged compared to control tissues in both experiments (***Figure 2a, 2b, right panels, Supplemental Figure 6***). Animals co-gavaged with ARA/EPA revealed an ARA distribution pattern similar to control or EPA alone. This suggests that EPA may compete with ARA, at least in part, by limiting ARA tissue accumulation.

**Figure 2.**
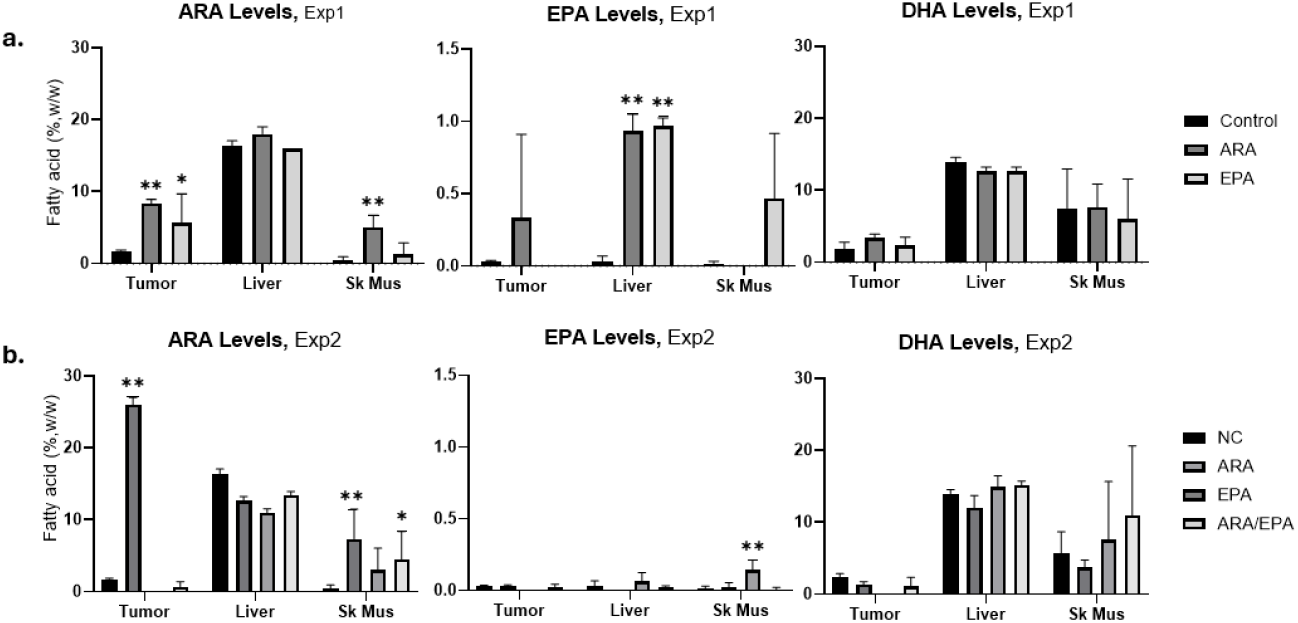
ARA, EPA and DHA levels in Tumor, Liver, and SK Mus (Skeletal Muscle) of control and gavage mice in a) Exp1 and b) Exp2. Significance was calculated by two-way ANOVA followed by Tukey’s multiple comparison test; *p<0.05 and **p<0.01.

### 3.6. DHA and D-DHA dosing suppressed liver ARA

Surprisingly, D-DHA appeared to support higher skeletal muscle ARA (***Figure 3a, left panel, Supplemental Figure 7***), EPA levels in DHA and D-DHA gavaged animals rose to 1-2%, suggesting either retroconversion of DHA to EPA or accumulation of EPA not converted to DHA [29] (***Figure 3a, right panel, Supplemental Figure 7***). Skeletal muscle EPA increased to about 0.5%, although this increase was not significant (***Figure 3a, right panel, Supplemental Figure 7)***. In response to DHA gavage, liver DHA rose to 20% from very low levels in controls and about 20% in D-DHA-dosed animals (***Figure 3b, Supplemental Figure 7***). Despite the rise in EPA with DHA dosing, we observed no D-EPA, which would have indicated retroconversion of D-DHA to D-EPA. These results suggest a strong correlation between elevated ω3 HUFA levels and the complete inhibition of tumor growth, positioning DHA and D-DHA as potent candidates for future anti-cancer therapies.

**Figure 3.**
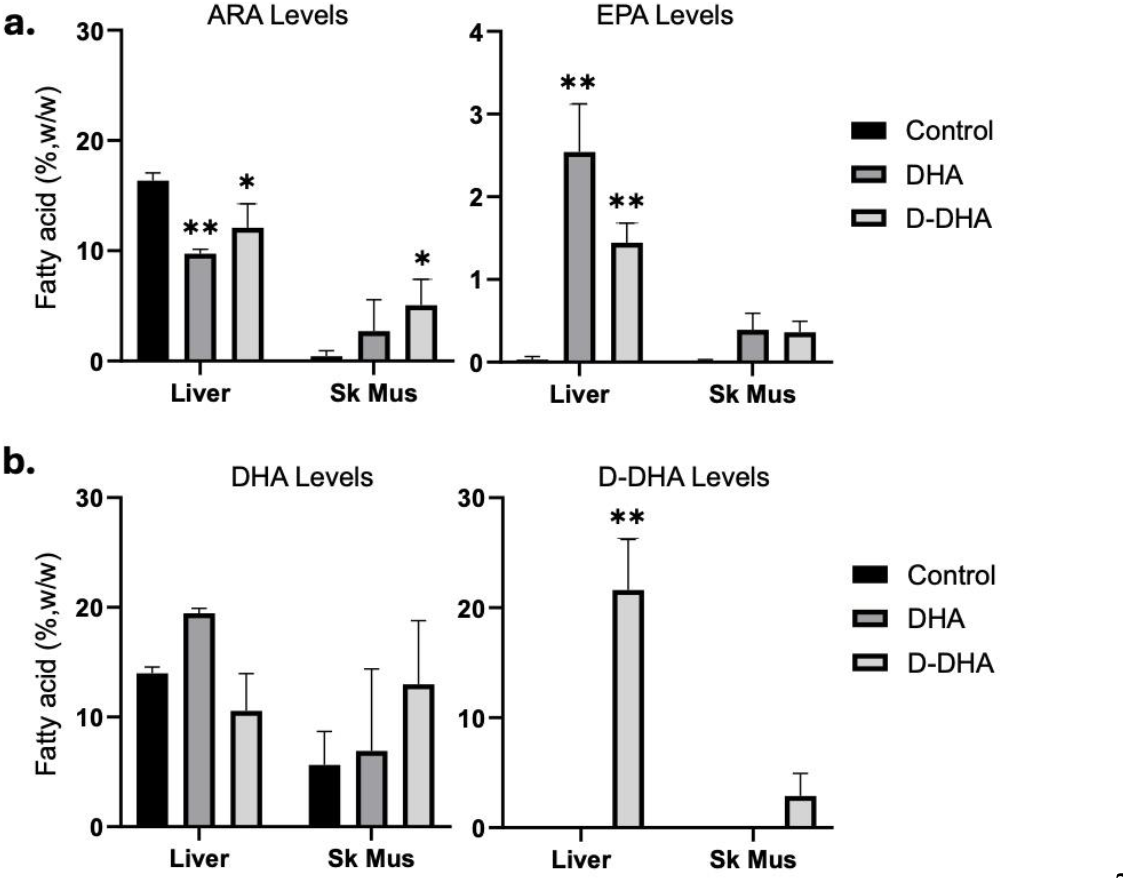
ARA, EPA, DHA, and D-DHA levels in Liver, and SK Mus (Skeletal Muscle) of control, DHA, and D-DHA gavage mice in Exp 2. a) ARA and EPA in Liver and SK Mus (Skeletal Muscle). DHA and D-DHA modulated ARA and EPA levels in both tissues. b) Neither DHA nor D-DHA gavage altered their respective levels in either tissue; D-DHA was only present in D-DHA gavaged mice. Significance differences from control was calculated by two-way ANOVA followed by Tukey’s multiple comparison tests; *p<0.05 and **p<0.01.

### 3.7. HUFA biosynthetic gene expression

The expression levels of key enzymes involved in unsaturated fatty acid metabolism (*Fads2, Elovl5, Fads1, Elovl2*; ***Supplemental Figure 8***) were significantly altered in the liver and tumor tissues of the HUFA-gavaged mice. While EPA-gavaged mice showed unaltered expression in tumors or skeletal muscle, they displayed significantly increased *Fads1* expression in the liver. This is notable as EPA is modestly yet significantly increased in livers of EPA-gavaged mice (***Supplemental Figure 9a, 9b, 9c***), consistent with *Fads1* production of EPA from its 20:4ω3 precursor, eicosatetraenoic acid (***Supplemental Figure 8***). A hepatic positive feedback loop for liver EPA generation when dietary EPA levels are high would explain these results. DHA and D-DHA significantly reduced the expression of *Fads1* in the liver and increased *Fads1* levels in skeletal muscle (***Supplemental Figure 9d, 9e***). The *Fads2* expression patterns were less distinct, only showing elevated *Fads2* in ARA- and D-DHA-gavaged livers (***Supplemental Figure10***). The elevated *Fads2* levels in ARA-driven tumors warrant future investigation of DPA (22:5ω6) with contributory pro-tumorigenic function.

*Elovl2*, the condensing enzyme in the first step of carbon chain elongation length of both ARA and EPA derivatives, was significantly elevated in tumors from EPA-gavaged mice. Elovl2 levels were unchanged regardless of HUFA exposure (***Supplemental Figure 11a, 11b, 11c, 11e***), except for a substantial reduction in DHA- and D-DHA-gavaged livers (***Supplemental Figure 11d***). In contrast, tumor levels of *Elovl5* were significantly increased in response to ARA-gavage (***Supplemental Figure 12a***). In the liver, levels of *Elovl5* were unchanged by DHA/D-DHA exposure but significantly reduced in response to EPA gavage (***Supplemental Figure 12b, 12d***). *Elovl5* levels were not significantly affected by HUFAs in skeletal muscle (***Supplemental Figure 11c, 11e***). These metabolic shifts reflect the complex interplay between fatty acid administration and enzymatic regulation, suggesting a deeper relationship between high-dose dietary HUFA fatty acids, tumor biology, and HUFA metabolism enzyme expression patterns.

## 4. Discussion

The current SoC for high-risk NB is typically divided into three phases: *induction* (chemotherapy; cisplatin, etoposide alternating with vincristine, cyclophosphamide, and doxorubicin; tumor resection), *consolidation* (myeloablative chemotherapy, stem cell transplant, radiation), and *maintenance* (high-dose isotretinoin treatment and immunotherapy). The use of multiple chemotherapeutic agents during the induction and consolidation phases of high-risk therapy is genotoxic to pediatric patients. Survivors often encounter lifelong health problems, including cognitive dysfunction, major joint replacements, and multiple organ dysfunction. They also have a 14% chance of developing a secondary malignancy over the ten years following chemotherapy [30,31]. Efforts to reduce the need or dosage requirements of genotoxic compounds are of significant continued importance. Reduced dosages of the genotoxic SoC components may improve overall health and survivorship in NB patients and influence SoC in other childhood cancers. The results reported here provide an attractive alternative for consideration. In addition to the successful use of ultra-high dose HUFA to treat traumatic brain injury, hypertriglyceridemia, and hypertension [16–18], we have demonstrated that ultra-high dose EPA and DHA can completely abrogate tumor formation in a syngeneic model of pediatric NB. In those previous efforts, tolerance of HUFA doses was excellent, and typically limited to occasional fishy burps and rarely GI discomfort. No major metabolic disturbances, including unacceptably increased bleeding/hemorrhaging, were observed. Our results have direct implications for prophylactic applications, use against metastatic disease, and therapeutic intervention.

The mechanism of action of EPA and DHA in blocking tumors will likely be via multiple underlying processes. Both ω3 HUFAs compete with ARA for incorporation into some, but not all, membranes, from which they are liberated before conversion to signaling molecules, thus reducing the amount of ARA available for this purpose. Once liberated, typically by a phospholipase A2, they compete with ARA for access to the COXs, LOXs, and other enzymes for conversion to prostaglandins, leukotrienes, thromboxane, and related molecules, which signal for a wide range of functions, including inflammation, thrombosis, vessel tone, and chemotaxis. Many studies also report their conversion to specialized pro-resolving lipid mediators [7]. The capillary endothelium is usually rich in ARA, suggesting that angiogenesis would be enhanced in ample ARA and inhibited when ARA is limiting [32].

This total suppression of tumor development further highlights the powerful anti-oncogenic effects of high-dose ω3 HUFAs like EPA and DHA. DHA, which forms nascent peroxyl radicals in response to ROS [33], contributes to the lipid hydroperoxide chain reaction. D-DHA, which has its bis-allylic hydrogens replaced with deuterium, is highly refractory to ROS-mediated damage *in vivo* [20,34]. Our results suggest an inherent value of an *in vivo* approach to prophylactic or therapeutic studies involving HUFAs. Previous *in vitro* studies using HUFAs on cell lines frequently report ROS-based cellular damage is a primary mechanism of cell death [15,35–37]. HUFAs are indeed sensitive to oxidation, and in 21% atmospheric oxygen of *in vitro* cell line studies, HUFA oxidation is robust

[37,38]. In our *in vivo* study, however, where oxidation rates of HUFA are already substantially less due to single-digit tissue oxygen levels *in vivo* [39,40], ROS-resistant D-DHA resulted in complete tumor inhibition, equivalent to ROS-sensitive EPA and DHA, suggesting that a non-ROS mechanism may be a primary driver. Future *in vivo* studies will elucidate the non-ROS mechanisms of DHA- and EPA-mediated tumor inhibition.

Our experiments showed that ARA promotes higher tumor incidence and earlier latency than other groups, suggesting that a pro-inflammatory environment derived from ARA derivatives favors cancer cell growth by releasing chemotaxis molecules. Increasing the ARA dose in Exp-2 did not result in more tumors but led to our observation of enhanced apparent angiogenesis around the tumor area, which was not seen with a lower dose of ARA. We posit that the ARA mechanism of action has both angiogenic and tumorigenic effects.

EPA/DHA doses higher than 20 g/d have been used on a sustained basis (months to years) in humans for the treatment of traumatic brain injury [17], 12 g/d to 15 g/d in hypertriglyceridemia [18] or hypertensive patients [16], and in at least one pregnant woman who gave birth successfully [41], and all doses were well tolerated. Based on observations in individuals with high circulating EPA levels, enhanced bleeding times might be expected, though a recent meta-analysis showed mild effects up to 4 g/d [42]. Our doses of DHA and EPA on a human calorie-equivalent basis are under 15 g/d (7.9 to 14.6 g/d, **Table 1**). The DHA level that was effective at suppressing all tumorigenesis is 12.2 g/d (human dose equivalent). This is about 3-fold the normal chronic ω3 EPA+DHA dose of prescription Lovaza™ used chronically to treat hypertriglyceridemia.

## 5. Conclusions

These results suggest that HUFA use may have prophylactic efficacy and therapeutic use targeting primary tumor growth and metastatic proliferation. With ω6 ARA enhancing and ω3 EPA/DHA/D-DHA blocking, their collective mechanisms that influence tumorigenesis may be directly opposed, where ω3 EPA and DHA shuts off anomalous, excess pro-inflammatory and proliferative signaling that can be enhanced by ω6 ARA. This relationship may apply more widely than NB and *MYCN*-driven/implicated cancers. Further, the high patient tolerance for HUFA without major deleterious side effects offers an attractive alternative to genotoxic chemotherapy and radiation inherent to current SoC, thus warranting further pre-clinical and clinical investigation.

## Supplementary Materials

The following supporting information can be downloaded at www.mdpi.com/xxx/s1. Figure S1: Gas chromatograms of fatty acids derived from the oils used for oral gavage; Figure S2: Quantitative profile of the oils used for dosing; Figure S3: Fatty acid composition of base chow; Figure S4: Fatty acid profile of Neuro 2a cells; Figure S5: Syngeneic CDX model of *MYCN*-driven neuroblastoma; Figure S6: Synthesis pathways of ω6 and ω3 HUFAs; Figure S7: Expression of *Fads1* in selected tissues; Figure S8: Expression of *Fads2* in selected tissues; Figure S9: Expression of *Elovl2* in selected tissues; Figure S10: Expression of *Elovl5* in selected tissues. Supporting Data: Mass spectrometry results for HUFA analysis (in Excel files)

## Author Contributions

Conceptualization, V.P., Y.N.L., J.T.B. and J.T.P.; Methodology, V.P., Y.N.L., L-R.E.B., H.G.P., J.T.B. and J.T.P.; Validation, V.P., Y.N.L., J.T.B. and J.T.P.; Formal analysis, V.P., Y.N.L., H.G.P., J.T.B. and J.T.P.; Investigation, V.P., Y.N.L., H.G.P., L-R.E.B., M.R., J.T.B. and J.T.P.; Resources, J.T.B. and J.T.P.; Data curation, J.T.B. and J.T.P.; Writing-original draft, V.P., L-R.E.B., J.T.B. and J.T.P.; Writing-review & editing, V.P., J.T.B. and J.T.P.; Visualization, V.P., J.T.B. and J.T.P.; Supervision, H.G.P., J.T.B. and J.T.P.; Project administration, J.T.B. and J.T.P.; Funding acquisition, J.T.P. All authors have read and agreed to the published version of the manuscript.

## Funding

This study was supported by the Cancer Prevention and Research Institute of Texas Grant RR180034 (PI J.T.P) and by a gift from Lake Austin Marina

### Institutional Review Board Statement

Not applicable.

### Informed Consent Statement

Not applicable.

## Data Availability Statement

The original contributions presented in this study are included in the article/supplementary material. Further inquiries can be directed to the corresponding author(s).

## Acknowledgments

Not applicable.

