## Supplemental Material for "Ultra-high dose oral *ω*3 docosahexaenoic acid (DHA) or eicosapentaenoic acid (EPA) block tumorigenesis in a *MYCN*-driven neuroblastoma model"

### *Supplemental Figure 1.*

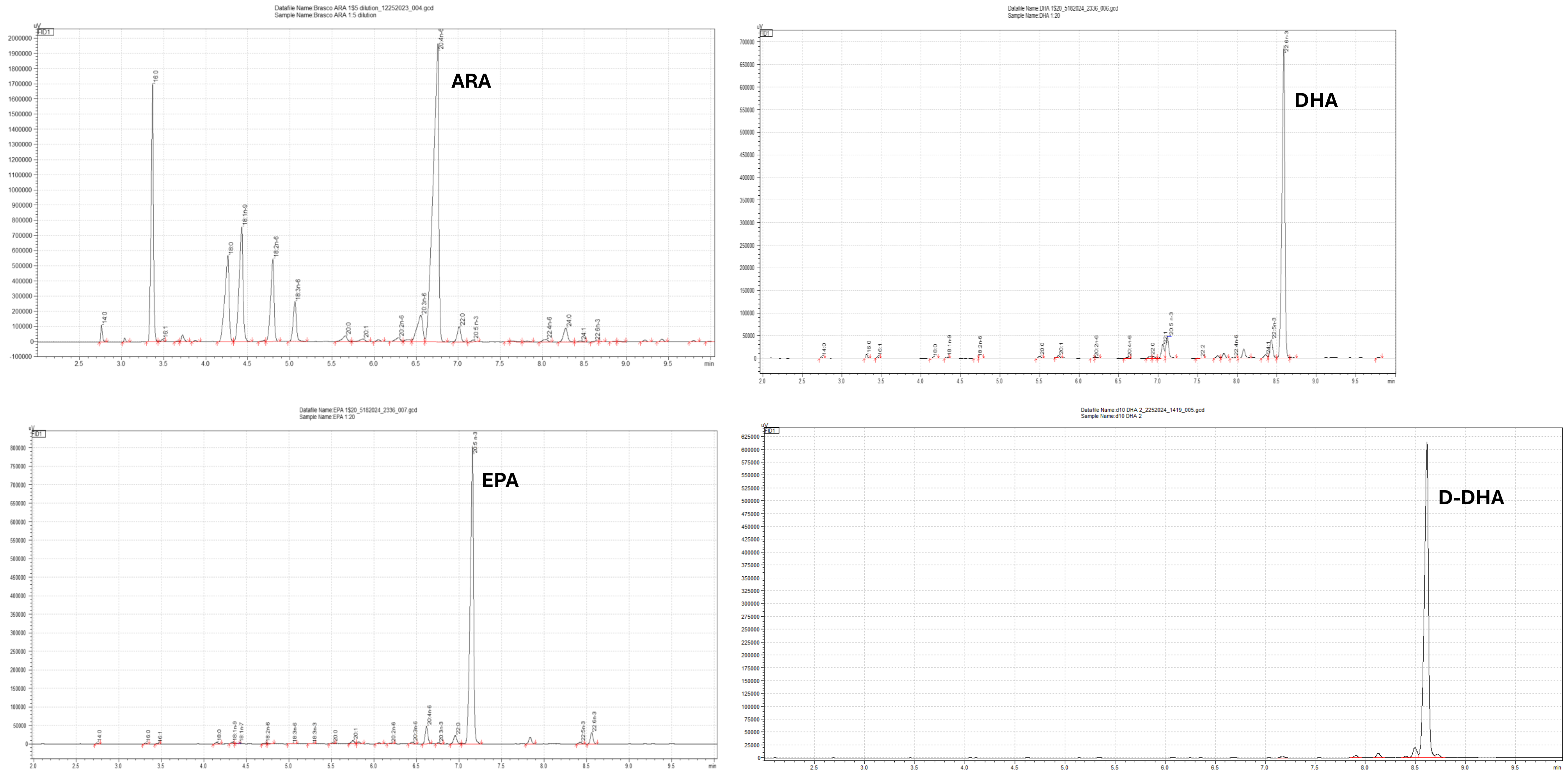

**Supplemental Figure 1.** Gas chromatograms of the oils used for oral gavage.

*Supplemental Figure 2.*

| Dosing Oils | | | |
|---|---|---|---|
| | ARA | EPA | DHA |
| 14:00 | 0.69% | 0.13% | 0.18% |
| 16:00 | 14.94% | 0.14% | 0.51% |
| 16:01 | 0.13% | 0.05% | 0.17% |
| 18:00 | 9.26% | 0.33% | 0.08% |
| 18:1n-9 | 11.02% | 0.31% | 0.17% |
| 18:2n-6 | 7.20% | 0.12% | 0.06% |
| 20:00 | 0.73% | 0.19% | 0.42% |
| 20:01 | 0.35% | 0.73% | 0.47% |
| 20:2n-6 | 0.49% | 0.17% | 0.08% |
| **20:4n-6 (ARA)** | **44.19%** | **3.94%** | **0.20%** |
| 22:00 | 1.41% | 2.27% | 0.56% |
| 22:01 | 0.00% | 0.00% | 3.01% |
| **20:5n-3 (EPA)** | **0.17%** | **87.64%** | **5.65%** |
| 22:02 | 0.00% | 0.00% | 0.14% |
| 22:4n-6 | 0.32% | 0.00% | 0.31% |
| 24:00:00 | 1.58% | 0.00% | 0.00% |
| 24:01:00 | 0.10% | 0.00% | 0.87% |
| 22:5n-3 | 0.00% | 0.41% | 4.63% |
| **22:6n-3 (DHA)** | **0.04%** | **2.67%** | **82.46%** |
| **Σω3 HUFA** | 0.21% | 90.72% | 92.74% |
| **Σω6 HUFA** | 44.51% | 3.94% | 0.51% |

***Supplemental Figure 2.*** Compositional analysis of the oils used for dosing presented as a quantitative fatty acid profile.

*Supplemental Figure 3.*

| Base Chow analysis | | |
|---|---|---|
| | **RT** | **Area** |
| 10:0 | No peak is detected. | 0.00% |
| 12:0 | 2.178 | 0.03% |
| 12:1 | 2.344 | 4.92% |
| 14:0 | 2.739 | 0.64% |
| 14:1 | 2.849 | 4.34% |
| 16:0 | 3.32 | 16.39% |
| 16:1 | 3.448 | 0.81% |
| 18:0 | 4.152 | 2.83% |
| 18:1n-9 | 4.344 | 12.65% |
| 18:1n-7 | No peak is detected. | 0.00% |
| 18:2n-6 | 4.767 | 47.95% |
| 18:3n-6 | 4.944 | 0.03% |
| 18:3n-3 | 5.259 | 5.74% |
| 20:0 | 5.455 | 0.21% |
| 20:1 | 5.7 | 0.20% |
| 20:2n-6 | 6.153 | 0.07% |
| 20:3n-6 | No peak is detected. | 0.00% |
| 20:4n-6 | 6.584 | 0.13% |
| 20:3n-3 | No peak is detected. | 0.00% |
| 22:0 | 6.833 | 0.31% |
| 22:1 | 7.064 | 0.07% |
| 20:5n-3 | 7.1 | 0.61% |
| 22:2 | 7.466 | 0.16% |
| 22:4n-6 | 7.953 | 0.13% |
| 24:0 | 8.072 | 0.26% |
| 24:1 | 8.289 | 0.04% |
| 22:5n-3 | 8.407 | 0.18% |
| 22:6n-3 | 8.546 | 1.30% |
| Sum | | 100.00% |

***Supplement Figure 3*****:** Base chow analysis showing the fatty acid composition of the chow diet (%w/w)

***Supplemental Figure 4.***

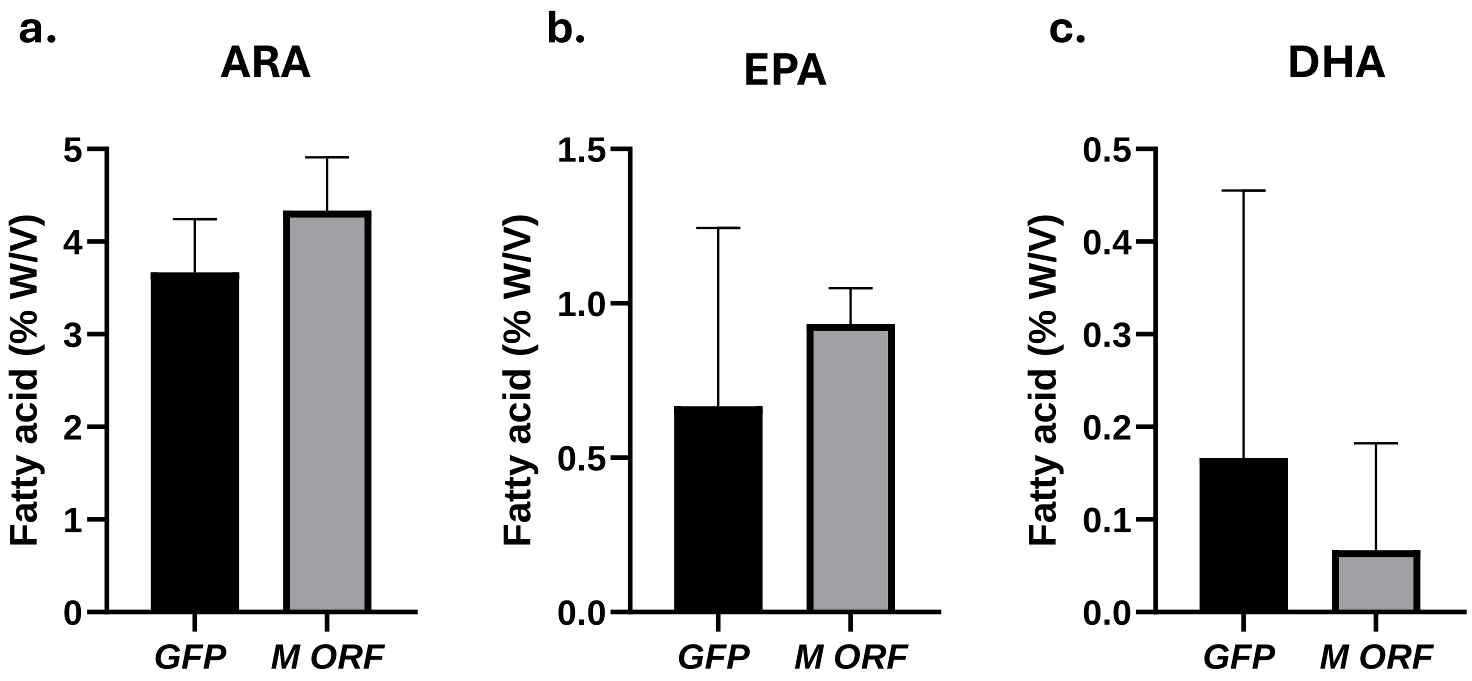

***Supplement Figure 4*: Fatty acid profile of Neuro 2a cells: a), b),** and **c)** Bar graphs represent ARA, EPA, and DHA levels in the Neuro 2a cells expressing MYCN compared to Control.

### *Supplemental Figure 5.*

**a.**

| **Neuro 2a (Experiment 1)** | | | |
|---|---|---|---|
| **Groups** | **Tumor Length (mm)** | **Tumor width (mm)** | **Tumor weight (g)** |
| *ARA* | 20 | 10 | 0.39 |
| | 9 | 10 | 0.29 |
| | 12 | 12 | 0.33 |
| | 11 | 10 | 0.24 |
| | 9 | 8 | 0.19 |
| *EPA* | 9 | 7 | 0.05 |
| | 4 | 7 | 0.06 |
| | 7 | 8 | 0.11 |

**b.**

| **Neuro 2a (Experiment 2)** | | | |
|---|---|---|---|
| **Groups** | **Tumor Length (mm)** | **Tumor width (mm)** | **Tumor weight (gm)** |
| Control | 3 | 3 | 0.10 |
| | 3 | 3.4 | 0.12 |
| | 2.5 | 3 | 0.12 |
| *Arachidonic acid ARA* | 7 | 5 | 0.13 |
| | 4.5 | 5 | 0.17 |
| | 5 | 4 | 0.19 |
| | 3 | 4 | 0.12 |
| | 6 | 5 | 0.20 |
| | 4 | 4 | 0.11 |
| *ARA+ EPA* | 4 | 5 | 0.09 |
| | 4 | 3.5 | 0.11 |
| | 3 | 4 | 0.08 |

**c.** **d.**

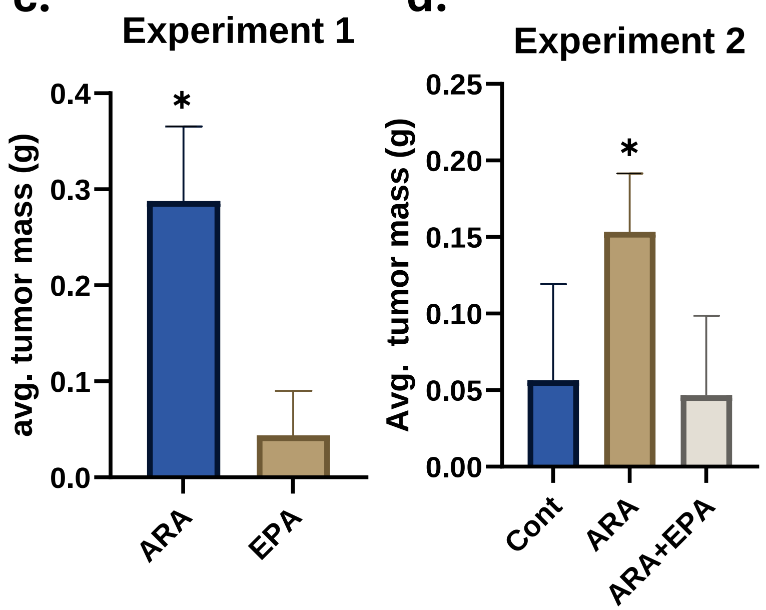

***Supplement Figure 5:*** **Syngeneic cell-derived xenograft (CDX) model of *MYCN*-driven neuroblastoma:** ARA EPA and ARA+ EPA were administered orally every 48 hours to the mice for seven days before cell injection. 2.5*10^6 *MYCN expressing* Neuro-2a cells were injected subcutaneously into the mice. ARA, EPA, and ARA+EPA feeding continued until the study endpoint in both experiment rounds. **a), b)** Summary of tumor size and mass with ARA and EPA in Exp1 and control, ARA and ARA+EPA in Exp2, respectively. **c), d)** Bar graph depicting the change in average tumor mass across different experimental groups, showing the highest tumor volume in the ARA-treated group in both experiment rounds. Significance was calculated using a t-test; *p<0.05 and **p<0.01.

### *Supplemental Figure 6.*

| **Figure 2 A** | **EXP 1** | **NC** | **NC** | **NC** | **ARA** | **ARA** | **ARA** | **EPA** | **EPA** | **EPA** |
|---|---|---|---|---|---|---|---|---|---|---|
| **ARA** | Tumor | 1.62 | 1.883 | 1.78 | 9 | 8 | 8 | 2 | 10 | 5 |
| | Liver | 15.937 | 16 | 17.156 | 19 | 18 | 17 | 16 | 16 | 16 |
| | Sk Mus | 0 | 0.986 | 0.357 | 4 | 4 | 7 | 3 | 1 | 0 |
| **EPA** | Tumor | 1.62 | 1.883 | 1.78 | 9 | 8 | 8 | 2 | 10 | 5 |
| | Liver | 15.937 | 16 | 17.156 | 19 | 18 | 17 | 16 | 16 | 16 |
| | Sk Mus | 0 | 0.986 | 0.357 | 4 | 4 | 7 | 3 | 1 | 0 |
| **DHA** | Tumor | 1.8 | 2.776 | 0.77 | 3 | 4 | 3 | 1 | 3 | 3 |
| | Liver | 13.406 | 14.55 | 14.002 | 13 | 12 | 13 | 12 | 13 | 13 |
| | Sk Mus | 13 | 2.103 | 7.437 | 4 | 10 | 9 | 12 | 5 | 1 |

| **Figure 2 B** | **EXP 2** | **NC** | **NC** | **NC** | **ARA** | **ARA** | **ARA** | **EPA** | **EPA** | **EPA** | **ARA/EPA** | **ARA/EPA** | **ARA/EPA** |
|---|---|---|---|---|---|---|---|---|---|---|---|---|---|
| **ARA** | Tumor | 1.62 | 1.883 | 1.78 | 27 | 25 | 26 | 0 | 0 | 0 | 0.58 | 1.46 | 0 |
| | Liver | 15.937 | 16 | 17.156 | 13 | 12 | 13 | 11.555 | 10.937 | 10.446 | 13.958 | 12.943 | 13.365 |
| | Sk Mus | 0 | 0.986 | 0.357 | 12 | 6 | 4 | 0.668 | 6.447 | 1.919 | 3.883 | 8.669 | 0.928 |
| **EPA** | Tumor | 0.04 | 0.025 | 0.02 | 0.02 | 0.03 | 0.04 | 0 | 0 | 0 | 0.03 | 0.04 | 0 |
| | Liver | 0 | 0.07 | 0.029 | 0 | 0 | 0 | 0 | 0.101 | 0.1 | 0.032 | 0.018 | 0.03 |
| | Sk Mus | 0 | 0.017 | 0.032 | 0 | 0.06 | 0 | 0.2 | 0.07 | 0.168 | 0 | 0 | 0.026 |
| **DHA** | Tumor | 1.8 | 2.776 | 2.51 | 1.16 | 1.83 | 1.12 | 0 | 0 | 0 | 0.97 | 2.44 | 0 |
| | Liver | 13.406 | 14.55 | 14.002 | 13 | 13 | 10 | 14.063 | 16.737 | 13.834 | 14.98 | 15.798 | 14.686 |
| | Sk Mus | 7.37 | 2.103 | 7.437 | 4.288 | 4.36 | 2.58 | 1.547 | 16.794 | 4.277 | 9.352 | 21.331 | 2.299 |

**Supplemental Table 1. Fatty Acid profiling:** ARA, EPA, and ARA/EPA were administered orally every 48 hours to the mice for seven days before cell injection. 2.5*10^6 *MYCN expressing* Neuro-2a cells were injected subcutaneously into the mice. ARA, EPA, and ARA/EPA feeding continued until the study's endpoint. The table represents HUFA levels in Tumor, Liver, and SK Mus (Skeletal Muscle) of control and gavage mice of Exp1 and Exp2, respectively.

### *Supplemental Figure 7.*

| Figure 3 | EXP2 | Control | Control | Control | DHA | DHA | DHA | D-DHA | D-DHA | D-DHA |
|---|---|---|---|---|---|---|---|---|---|---|
| ARA | Liver | 15.937 | 16 | 17.156 | 9.64 | 10.18 | 9.42 | 11.76 | 10.11 | 14.4 |
| | Sk Mus | 0 | 0.986 | 0.357 | 1 | 1.17 | 5.99 | 3.95 | 7.77 | 3.48 |
| EPA | Liver | 0 | 0.07 | 0.029 | 2.09 | 2.33 | 3.2 | 1.17 | 1.56 | 1.6 |
| | Sk Mus | 0 | 0.017 | 0.032 | 0.26 | 0.3 | 0.62 | 0.37 | 0.23 | 0.49 |
| DHA | Liver | 13.406 | 14.55 | 14.002 | 19.03 | 19.44 | 19.91 | 7.31 | 10.27 | 14.1 |
| | Sk Mus | 7.37 | 2.103 | 7.437 | 2.32 | 2.89 | 15.55 | 9.68 | 19.7 | 9.56 |
| D-DHA | Liver | 0 | 0 | 0 | 0 | 0 | 0 | 22.93 | 25.4 | 16.5 |
| | Sk Mus | 0 | 0 | 0 | 0 | 0 | 0 | 3.22 | 4.74 | 0.71 |

**Supplemental Table 2. Syngeneic cell-derived xenograft (CDX) model of *MYCN*-driven neuroblastoma:** DHA and D-DHA were administered orally every 48 hours to the mice for seven days before cell injection. 2.0*10^6 *MYCN expressing* Neuro-2a cells were injected subcutaneously into the mice. DHA and D-DHA feeding continued until the study endpoint. The table represents ARA, EPA, and DHA levels in Liver and SK Mus (Skeletal Muscle) of control and gavage mice of Exp 2.

***Supplemental Figure 8.***

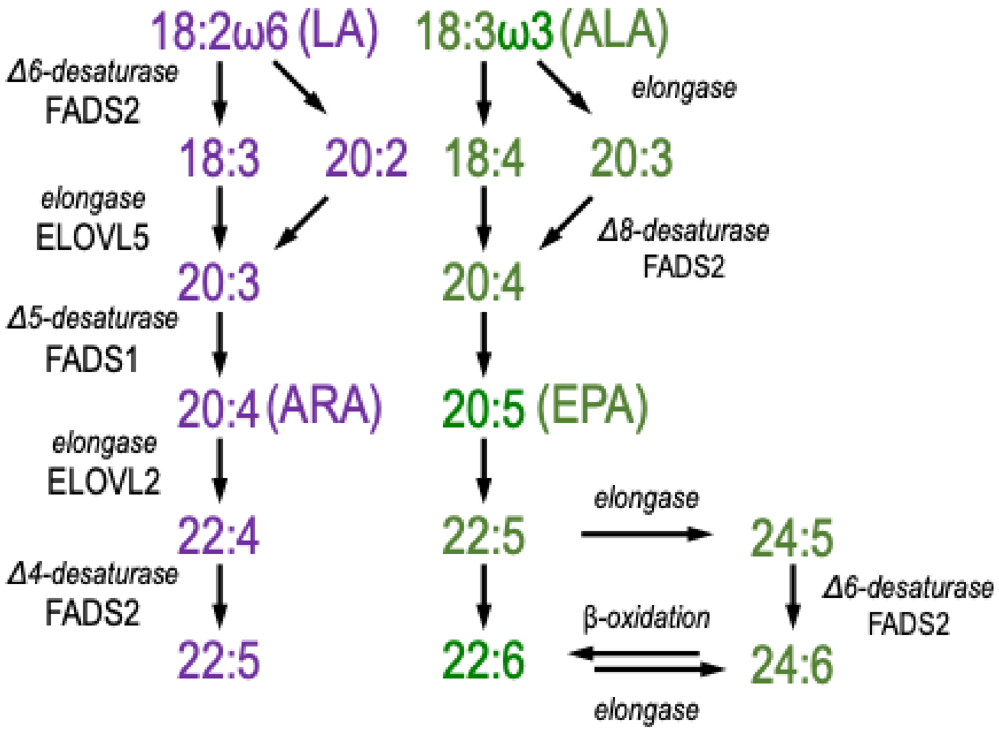

**Supplement Figure 6.** Synthesis pathways of ω6 and ω3 HUFAs

*Supplemental Figure 9.*

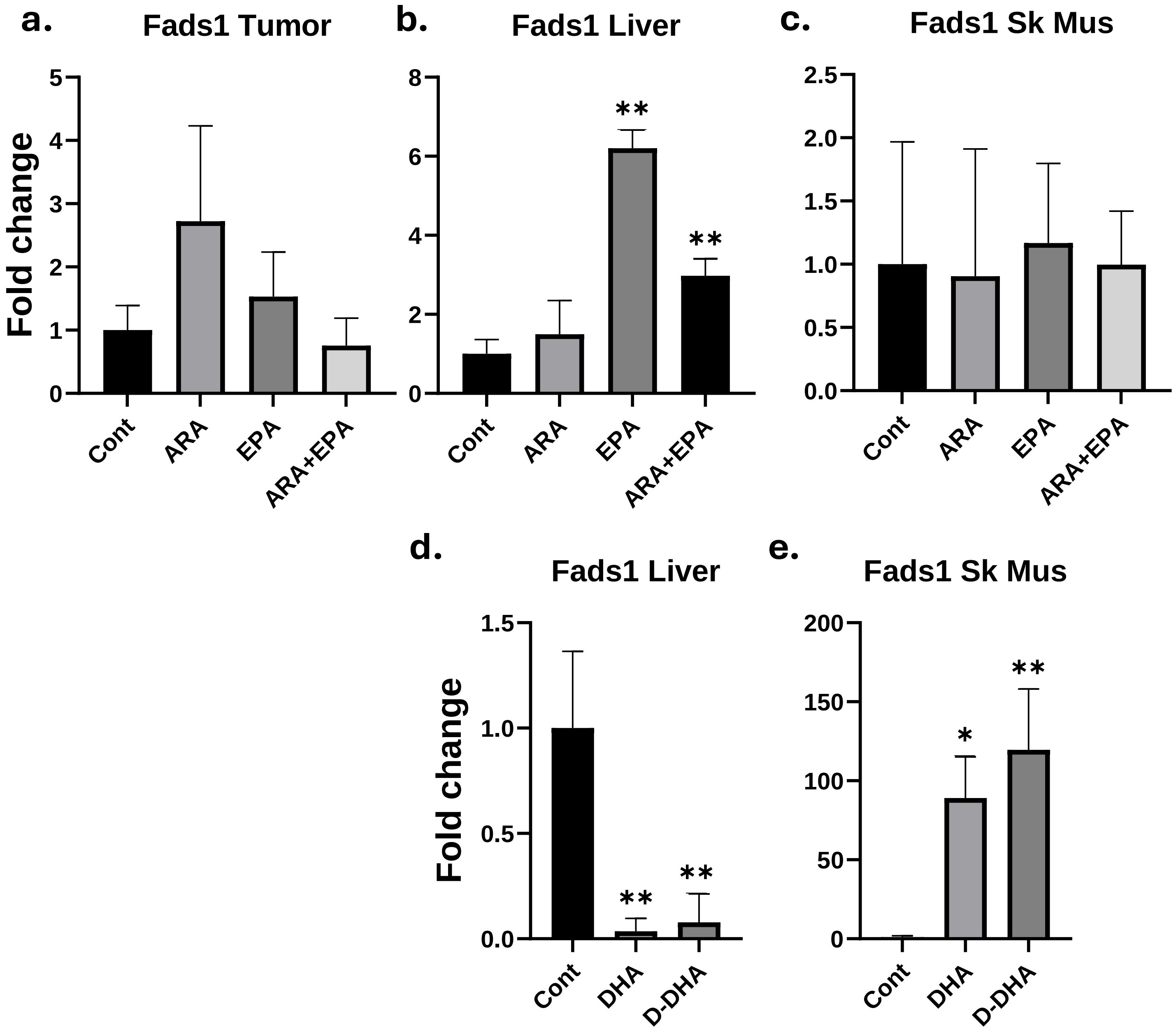

**Supplement Figure 7. Expression of *Fads1* in selected tissues: a), b), c), d), and e)** Bar graph represents *Fads1* levels in Tumor, Liver, and SK Mus (Skeletal Muscle) of control and gavage mice of Exp2; Significance was calculated using a Two-way ANOVA followed by Tukey's multiple comparison tests; *p<0.05 and **p<0.01.

*Supplemental Figure 10.*

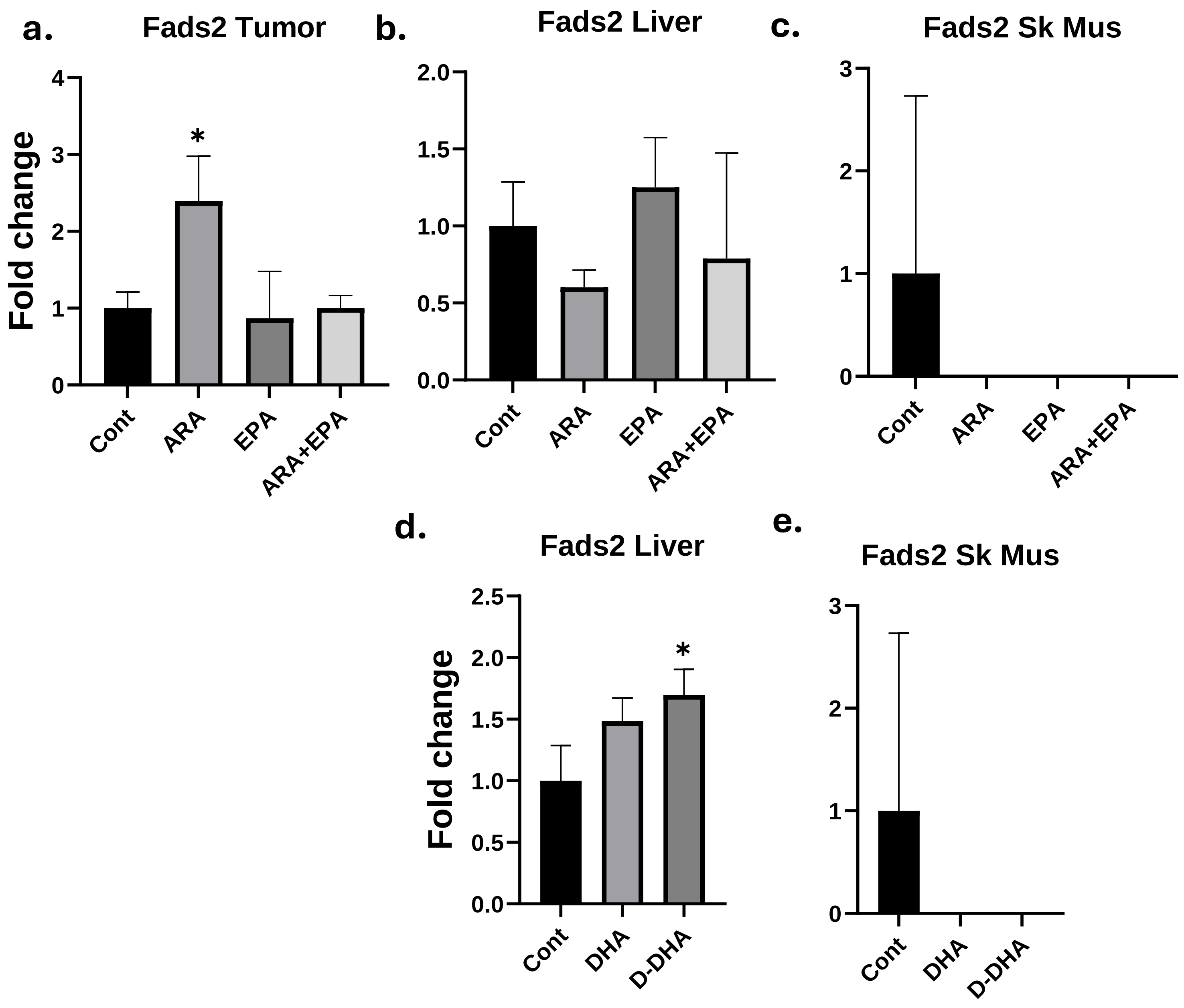

**Supplement Figure 8: Expression of *Fads2* in selected tissues: a), b), c), d), and e)** Bar graph represents *Fads2* levels in Tumor, Liver, and SK Mus (Skeletal Muscle) of control and gavage mice of Exp2; Significance was calculated using a Two-way ANOVA followed by Tukey's multiple comparison tests; *p<0.05 and **p<0.01.

*Supplemental Figure 11.*

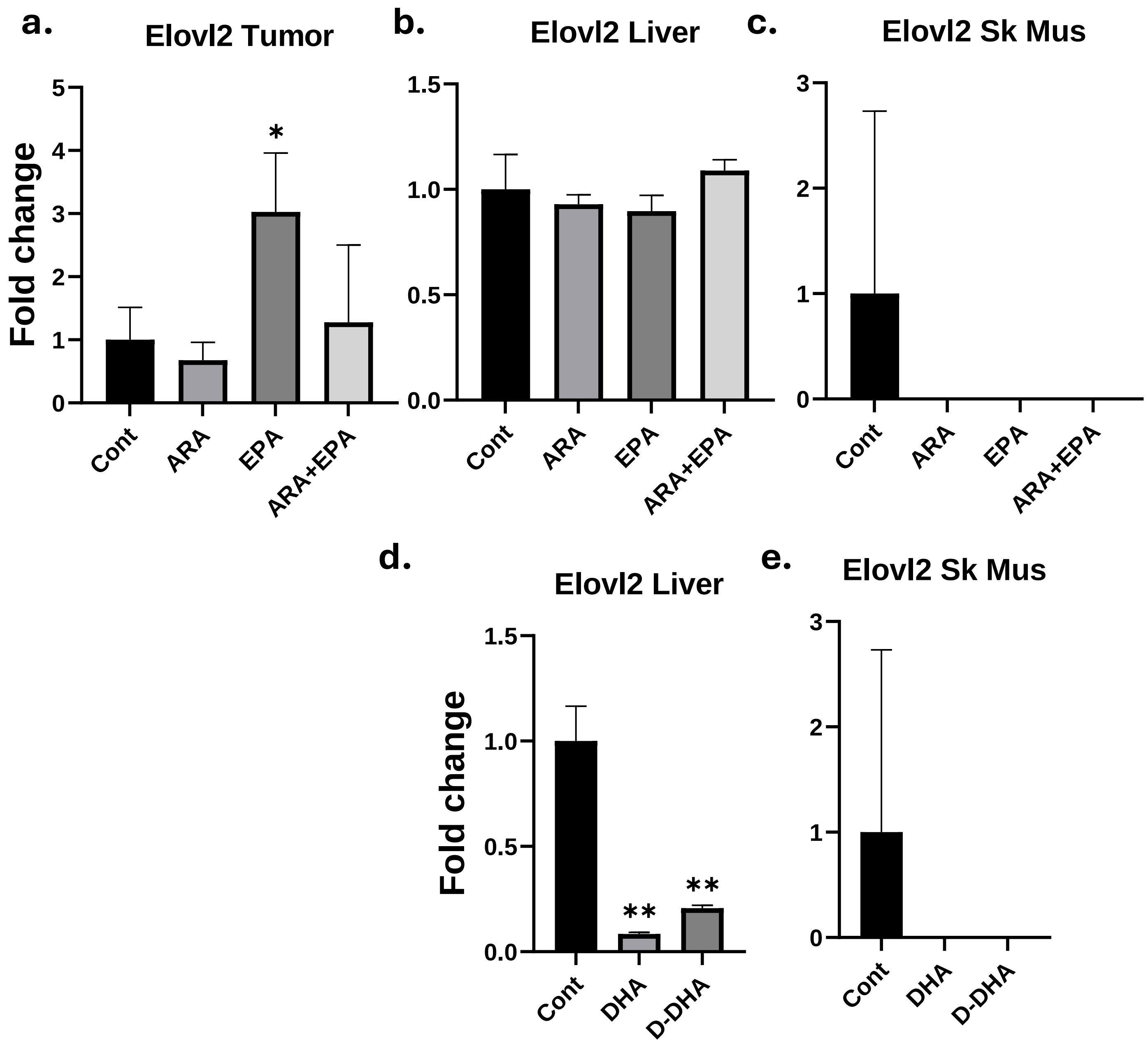

**Supplement Figure 8: Expression of *Elovl2* in selected tissues: a), b), c), d), and e)** Bar graph represents *Elovl2* levels in Tumor, Liver, and SK Mus (Skeletal Muscle) of control and gavage mice of Exp2; Significance was calculated using a Two-way ANOVA followed by Tukey's multiple comparison tests; *p<0.05 and **p<0.01.

### *Supplemental Figure 12.*

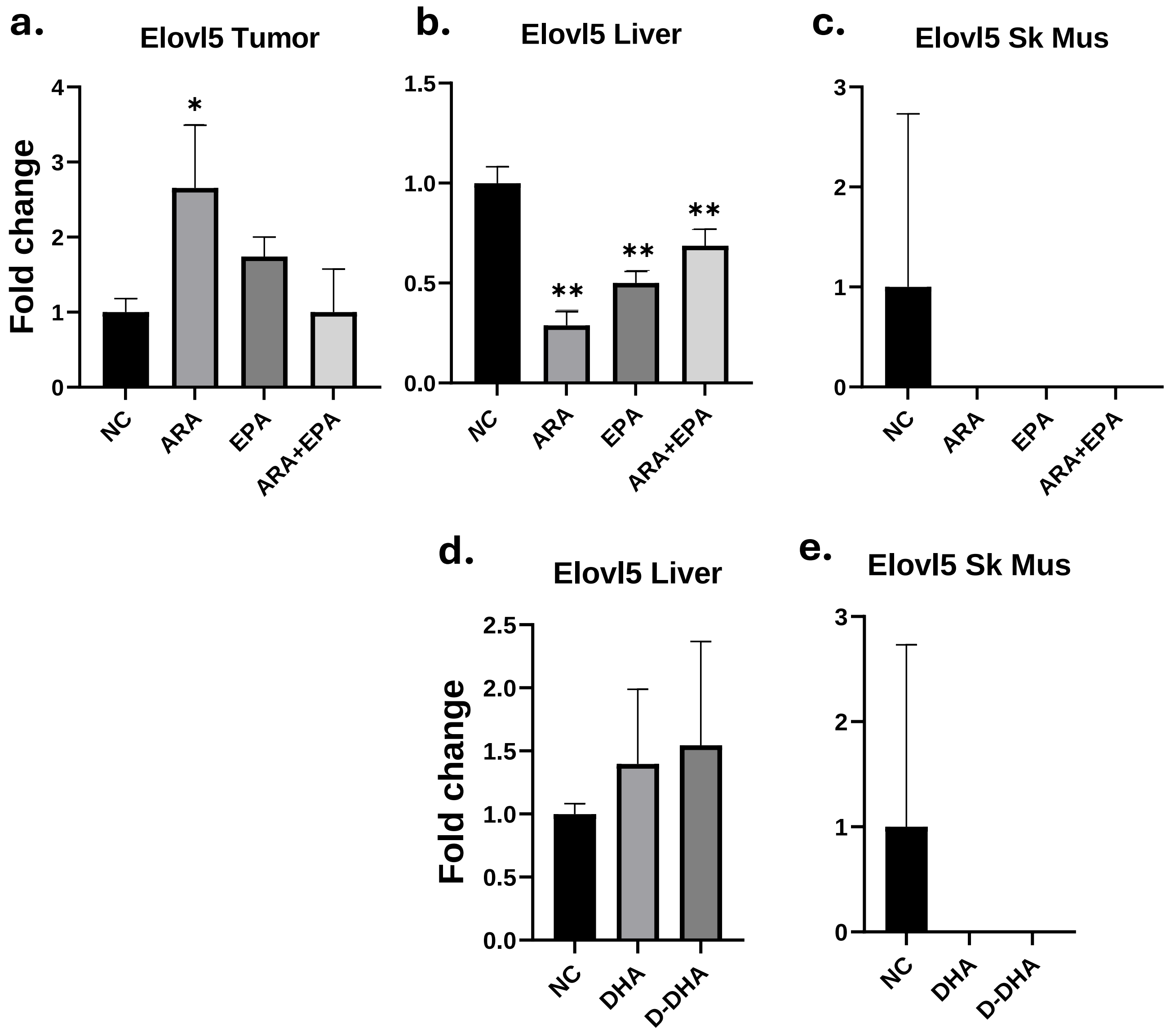

**Supplement Figure 10: Expression of *Elovl5* in selected tissues: a), b), c), d), and e)** Bar graph represents *Elovl5* levels in Tumor, Liver, and SK Mus (Skeletal Muscle) of control and gavage mice of Exp2; Significance was calculated using a Two-way ANOVA followed by Tukey's multiple comparison tests; *p<0.05 and **p<0.01.
